# A Transparent Ultrasound Array for Real-time Optical, Ultrasound and Photoacoustic Imaging

**DOI:** 10.1101/2021.11.09.467971

**Authors:** Haoyang Chen, Sumit Agrawal, Mohamed Osman, Josiah Minotto, Shubham Mirg, Jinyun Liu, Ajay Dangi, Quyen Tran, Thomas Jackson, Sri-Rajasekhar Kothapalli

## Abstract

**Objective and Impact Statement:** Simultaneous imaging of ultrasound and optical contrasts can help map structural, functional and molecular biomarkers inside living subjects with high spatial resolution. There is a need to develop a platform to facilitate this multimodal imaging capability to improve diagnostic sensitivity and specificity.

**Introduction:** Currently, combining ultrasound, photoacoustic and optical imaging modalities is challenging because conventional ultrasound transducer arrays are optically opaque. As a result, complex geometries are used to co-align both optical and ultrasound waves in the same field of view.

**Methods:** One elegant solution is to make the ultrasound transducer transparent to light. Here, we demonstrate a novel transparent ultrasound transducer (TUT) liner array fabricated using a transparent lithium niobate piezoelectric material for real-time multimodal imaging.

**Results:** The TUT array consisted of 64 elements and centered at ∼ 6 MHz frequency. We demonstrate a quad-mode ultrasound, Doppler ultrasound, photoacoustic and fluorescence imaging in real-time using the TUT array directly coupled to the tissue mimicking phantoms.

**Conclusion:** The TUT array successfully showed a multimodal imaging capability, and has potential applications in diagnosing cancer, neuro and vascular diseases, including image-guided endoscopy and wearable imaging.

## 1 Introduction

Ultrasound and optical imaging modalities are non-ionizing, portable, affordable and can be realized in various forms, from table top size to miniaturized endoscopes or wearable devices [1, 2]. Ultrasound (US) imaging provides the deep tissue structural information based on differences in acoustic impedance and complementary functional blood flow information through Doppler ultrasound [3]. Photoacoustic (PA) imaging, as a hybrid imaging modality maps optical absorption contrast of deep tissue with ultrasonic spatial resolution. For example, hemoglobin absorption based label-free imaging of vascular anatomy and functional oxygen saturation have been shown to be useful in diagnosing cancer, neurological, and vascular diseases [4–6]. Pure optical imaging methods such as fluorescence imaging enables biochemical information of targeted cells and tissue (e.g. autofluorescence from metabolic co-factors NAD/NADH: nicotinamide adenine dinucleotide) and therefore allows high diagnostic sensitivity and specificity [7–10]. Deep tissue US and PA imaging modalities provide higher imaging depth and better spatial resolution (scalable with ultrasound parameters, typically 1/100^*th*^ of an imaging depth, that is 0.5 mm spatial resolution at 5 cm depth) compared to deep tissue optical imaging with a spatial resolution of about 1/5^*th*^ of an imaging depth. However, when probing superficial depths (*<* 1 mm) optical imaging modalities provide relatively better spatial resolution (submicrons to a few microns). Therefore, a synergistic integration of optical, US and PA imaging technologies into a single multimodal imaging platform will provide complementary contrasts, penetration depths, and high spatial resolutions. They are desired in many biomedical applications to simultaneously image a set of structural, functional and molecular biomarkers.

Different combinations of optical and ultrasound imaging systems have been reported for different clinical applications. In cancer imaging, Fatakdawala et al., demonstrated *in vivo* imaging of oral cancer in a hamster model using a bench-top combination of fluorescence lifetime (FLI), PA and US imaging techniques [11]. FLI revealed biochemical (NADH) changes on the tissue surface, with a lower fluorescence lifetime for the oral cancer tissue compared to the surrounding tissue. US imaging provided underlying tissue morphology and microstructure, and PA imaging detected high vascularization within the cancerous tissue. Similarly, Tummers et al., performed multimodal US, PA and fluorescence imaging of a surgical removed pancreatic specimen obtained from a pancreatic ductal carcinoma (PADC) patient [12], who was intravenously administered with a near-infrared (NIR) fluorescent agent, Cetuximab-IRDye800, that binds to epidermal growth factor receptor. In this case, fluorescence imaging provided the surface projection of the targeted Cetuximab-IRDye800 agent, PA imaging showed the depth resolved optical absorption contrast from the IRDye800 and surrounding vasculature, and ultrasound imaging revealed the underlying tissue anatomy. For imaging atherosclerosis, a cardiovascular disease characterized by the accumulation of lipid plaques and several fibrous and cellular constituents, intravascular ultrasound (IVUS) and optical coherence tomography (OCT) technologies are commonly used in the clinics [13–15]. Recently, intravascular PA (IVPA) is also being actively studied for mapping deep tissue atherosclerosis based on high optical absorption contrast of plaque lipids in the NIR-IIb (1.5 µm – 1.7 µm) optical window [16–20]. Similarly, neuroscience studies also require high-resolution multiparametric hemodynamic information (cerebral blood flow, blood volume and oxygen saturation) obtained from optical and photoacoustic imaging for mapping resting state brain connectivity [21–23], studying neuromodulation [24], neurovascular coupling [25–27], and neuro diseases [28–30]. For this purpose, recently functional ultrasound (fUS) imaging, which provides high resolution images of microvascular blood flow, has been integrated with hemoglobin absorption-based PA vascular imaging [31].

However, the current experimental setups integrating fluorescence, US and PA technologies are limited to raster scanning the imaging device over the tissue sample, one imaging mode at a time [11]. Since real-time US imaging (e.g., IVUS and fUS) is performed using ultrasound transducer array, the most viable approach for real-time multimodal imaging is to integrate fluorescence (or other optical technologies) and PA imaging to the US imaging array based platform. However, optical opacity of conventional ultrasound transducers hinders co-axial and compact integration of the ultrasound transducer array with optical illumination and detection fibers. For example, real-time B-mode US and PA (USPA) imaging devices are developed by simply assembling optical fiber bundles around a conventional ultrasound transducer probe (Fig. 1a). Due to the physical separation between the two optical fiber bundles, optical illumination is not available below the surface of the ultrasound transducer up to 1 - 2 cm depth (see Figs. 1a-b) [32, 33]. To partially offset this problem and achieve co-aligned optical and ultrasound fields on the tissue surface, USPA devices are operated with long working distances (*>* 1 cm), visible as dark region (Fig. 1b), using water or ultrasound gel as the coupling medium between the tissue and the probe surface [34, 35]. This limits miniaturization of the devices and longitudinal *in vivo* imaging capabilities, introduces artifacts, increases ultrasound attenuation as well as ultrasound scattering if any bubbles are formed in the coupling medium. The requirement for long working distance also limits the imaging speed because of redundant data corresponding to non-illuminated region is also captured and processed by the data acquisition system.

**Figure 1:**
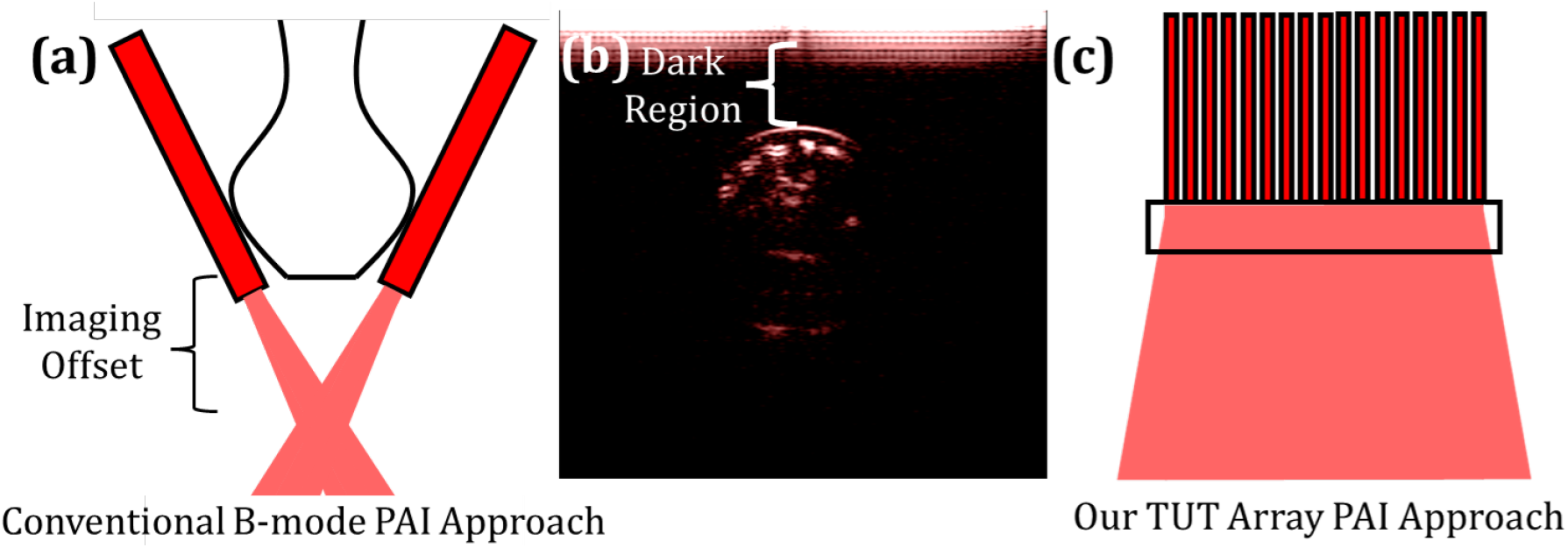
Photoacoustic tomography setup that employs a linear array. **(a)** Light needs to be attached from the sides, creating imaging offset.**(b)** A photoacoustic image of the finger shows the dark region that lacks the illumination. Acquired by Acoustic X. However, **(c)** a transparent linear array allows the light to be coupled from the backside, and removes the imaging offset.

The above challenges can be overcome by employing transparent ultrasound transducers (TUT) that allow light delivery through the transducer, as shown in Fig. 1c. By doing so, the ultrasound transducer becomes a part of the optical system, instead of an obstruction to the optics. This will not only significantly reduce the beam engineering challenges but will also lead to the development of a more compact, portable, wearable and versatile multimodal systems. For this purpose, both conventional piezoelectric materials [36–39] and capacitive micromachined ultrasound transducers (CMUTs) [40–42] have been studied for developing TUTs. Ilkhechi et al. reported transparent CMUT array for ultrasound imaging of a small size tissue phantom [41] and photoacoustic [42] imaging of pencil leads submerged in oil tank. These transparent CMUTs have not yet shown deep tissue US and real-time dual-modality USPA imaging capabilities. Further, CMUTs in general employ complex clean room fabrication processes, large bias voltages, and custom-developed integrated circuits for operation leading to their incompatibility with current clinical ultrasound systems [5]. All-optical photoacoustic detectors such as transparent optical ring resonators[43], and Fabry–P’erot etalons[44] have the ability to transmit light through them and into the tissue. However, these systems require additional laser and optical detectors, yet to be realized in the form of 1D or 2D arrays, and are not compatible with commercial ultrasound machines[45]. Moreover, these detectors cannot be used for ultrasound excitation/imaging required for dual-modality USPA imaging applications. While prior studies demonstrated the potential of transparent lithium niobate (LN) based single element TUTs for high sensitivity PA imaging [36–39], TUT-arrays are required for real-time multimodal imaging.

To address above-mentioned limitations, in this work, we introduced the first bulk 1D TUT-array using a transparent LN piezoelectric material and demonstrated its feasibility for a real-time multimodal deep tissue imaging. We characterized the TUT array using electrical and acoustic methods. The TUT array enabled co-alignment of acoustic and light pathways with minimal acoustic coupling. Imaging of tissue mimicking phantoms validated a quad-mode US, PA, Doppler ultrasound, and fluorescence imaging capabilities of the TUT array for providing respective structural, functional, and molecular information of the tissue without introducing any shadow regions. In the future, TUT arrays can have broad biomedical applications such as compact multimodal endoscopy or wearable imaging applications and also for photomediated ultrasound therapy for deep vein thrombosis or wound healing.

## 2 Results

### 2.1 TUT-Array Characterization

#### Array Design

A center frequency of 6.5 MHz was chosen to match commonly used diagnostic ultrasound devices. This can be achieved by a 0.5 mm thick LN. The element width of 0.2 mm was chosen to be less than 0.6×(element thickness) and greater than *λ/*2 to avoid spurious resonant modes. Here *λ* represents the ultrasound wavelength in tissue medium. When designing element pitch, it needs to be within the range from *λ/*2 to 3*λ/*2 to avoid grating lobes. Therefore, a pitch of 0.3 mm was chosen for a 6.5 MHz linear array [46] with a total of 64 elements and an element height of 5 mm.

#### Acoustic Stacking

The acoustic stacking of the transparent array is shown in Fig. 2a. Double side ITO coated LN was selected as the piezoelectric material due to its high optical transmission rate (> 80% in the NIR wavelengths) and good electromechanical coupling coefficient (49%). 64 elements were created by dicing 400 µm deep inside the 500 µm LN wafer, leaving 100 µm for shorting all elements as the common ground. A conductive glass slide was bonded to the LN and served as the first backing layer as well as the ground connection. An additional backing layer of transparent epoxy was placed on top of the glass slide to further reduce the acoustic reverberation. To individually address each element, a custom fabricated cable was anisotropic conductive film (ACF) bonded to the edge of the array as shown in Fig. 2a-b. To improve the ultrasound energy transmission, a quarter wavelength matching layer of Parylene-C (not shown in Fig. 2a schematic) was deposited for acoustic matching and waterproof layer.

**Figure 2:**
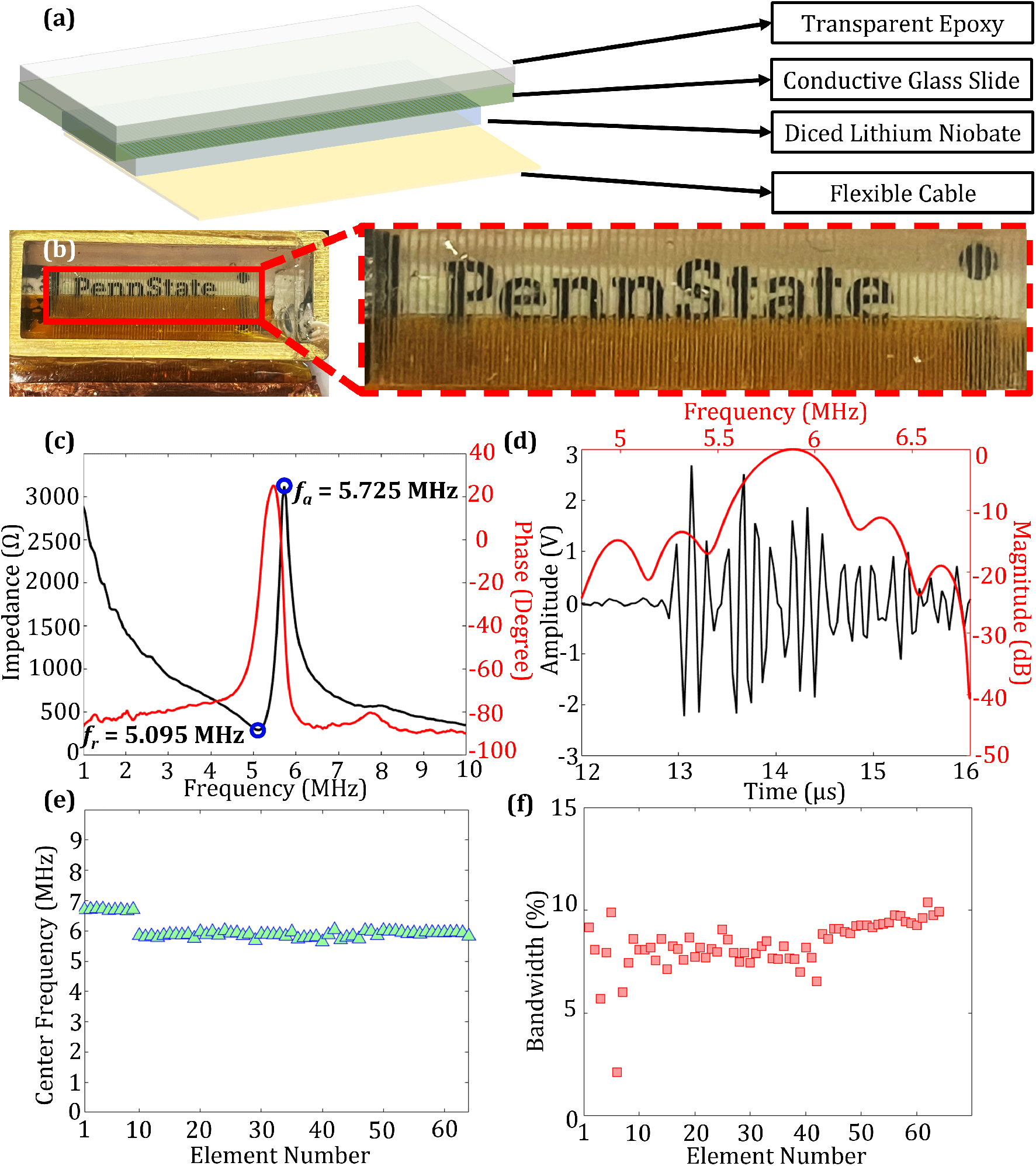
**(a)** Schematic of the transparent ultrasound transducer (TUT) array using double sided ITO coated transparent lithium niobate piezoelectric material. The matching layer of Parylene is not shown. **(b)** a photograph of the fabricated TUT linear array on top of the ”Penn State” sign. The TUT-array characterization results: **(c)** electrical impedance plots for element #32. The black line represents the impedance and the red line represents the phase **(d)** Two way pulse echo response for element #32. The black line represents the time domain signal and the red line represents the corresponding frequency response. The pulse echo characterization was conducted for all 64 elements. The determined **(e)** center frequencies and **(f)** bandwidths for each element are plotted.

The TUT-array design and step-wise fabrication is presented in the Methods section: TUT-Array fabrication and packaging. The design parameters for the TUT-array are summarized in the Table 1. Fig. 2a shows the schematic design and Fig. 2b shows the picture of the fabricated TUT linear array on top of a ”Penn State” logo. Although discontinuity was observed between the elements due to the dicing kerf, the letters are clearly readable throughout the TUT-array. Red dashed box shows that the gold traces from the flexible circuit were properly aligned and bonded to each LN element.

**Table 1:** Design Parameters for Transparent Linear Array.

| Parameters | Values |
| --- | --- |
| Center Frequency | 6.5 MHz |
| Number of Elements | 64 |
| Pitch | 0.3 mm |
| Element size T×H×W | 0.5 mm×5 mm×0.2 mm |
| Electrode Thickness (ITO) | 200 nm |
| Backing Thickness (Glass + Epoxy) | 10 mm |
| Matching Thickness (Parylene) | 80 $\mu\text{m}$ |
| T: thickness; H: height; W: width. ITO: indium tin oxide |  |

Following the fabrication, various tests to the TUT-array were conducted to determine its electrical and acoustic characteristics and their consistency across all 64 elements. The electrical impedance measurements were first conducted for each element of the TUT array and the characterization method was described in our literature [37]. A calibrated electrical impedance analyzer (Agilent E5100A, Keysight Technologies, Inc., Santa Rosa, CA, USA) was used to determine the phase and electrical impedance for each linear array element. Fig. 2c shows the measured input impedance and phase curve from the center element: #32. The resonance and anti-resonance frequency were found to be 5.095 MHz and 5.725 MHz, respectively. The electromechanical coupling coefficient was then calculated to be 0.456 according to the IEEE standard on piezoelectricity [47]. Pulse echo characterization was carried out by placing a flat metal target at ∼ 23 mm away from the TUT-array, while each element was excited with electrical pulses with 40 volts amplitude from the Vantage 256 (Vantage 256, Verasonics Inc., Kirkland, WA, USA). Fig. 2d shows typical pulse echo results obtained from the center element #32. The center frequency of the element was found to be 5.88 MHz, and -6 dB fractional bandwidth of 8.8%. This pulse echo procedure was swept for each element, and the measured center frequencies for all 64 elements were plotted in Fig. 2e and corresponding bandwidths were plotted in Fig. 2f. The first 9 elements showed a center frequency at 6.695 MHz ± 0.023 MHz, while the rest 55 elements exhibited a center frequency of 5.877 MHz ± 0.090 MHz. The difference can be attributed to the non-uniform pressing, which left an uneven sheet of bonding epoxy between the backing conductive glass slide and the LN. This was further confirmed by examining the bandwidths of the 64 elements in Fig. 2f, where the first 9 elements showed a lower bandwidth average and a larger standard deviation: the average fractional band-widths for first 9 elements was found to be 7.22% ± 2.35%, while the rest 55 elements showed an average fractional bandwidth of 8.49% ± 0.85%, but they are much lower compared to conventional medical ultrasound linear arrays in the range of 40% to 70% bandwidth.

### 2.2 Quad-mode Imaging Validation

Using the fabricated TUT arrays, we demonstrated a quad-mode US, PA, Doppler US and optical fluorescence imaging capabilities. **USPA Imaging**: The TUT-array was connected to the Vantage 256 ultrasound data acquisition system to perform real-time interleaved US and PA imaging (see Methods: Imaging system and data acquisition sequence) on a tissue phantom prepared using a solution mixture of agarose and silica powder. The phantom and imaging schematic is shown in Fig. 3a. The phantom consisted of four metal wire targets, each with a diameter of 50 *µ*m and dyed with India ink to generate strong photoacoustic contrast. Fig. 3a shows the approximate positions of the 4 photoacoustic targets along the imaging depth of the phantom, with approximately 5 mm distance between the targets. The tissue phantom also consisted of two ultrasound only targets (H1 and H2) in cylindrical shape filled with agar solution to mimic hypoechoic regions in the tissue medium (see Methods: Multimodal imaging phantom preparation). The TUT-array was directly placed on top of the phantom and the laser light irradiated the phantom through the TUT-array, demonstrating the advantage of using the TUT-array for dual-modality USPA imaging with minimal coupling. The corresponding US and PA imaging results of the phantom are demonstrated in Figs 3c-d. The US image in Fig. 3c clearly shows four micro metal wires (∼5 mm in axial plane and ∼3 mm in lateral plane) as these wire targets have different acoustic impedance compared to the background tissue mimicking medium. Due to limited bandwidth of the current TUT-array, the metal wire targets in the ultrasound images are not sharp (point like) in the depth axis. The gray shades on the lateral direction can be attributed to the cross-talk between the elements in the array with 80% dicing depth. As expected, the ultrasound contrast from hypoehoic targets appeared as dark regions (red circled) in Fig. 3c, with distinguishable boundary from the background ultrasound speckle contrast. The measured hypoehoic target diameter (∼5 mm) is closed to the real target size (4.75 mm). The PA imaging result in Fig. 3d showed depth-resolved optical absorption contrast from the four metal wires dyed with India ink. The locations of all four wires are clearly displayed at expected locations with sufficient PA contrast from the background. Considering the 50 µm diameter metal wire as the point target, we calculated the full-width half maxima from point spread function to quantify the lateral and axial resolution, and they were found to be 0.45 mm and 0.86 mm respectively for US and 0.8 mm and 1.1 mm respectively for PA at W3 in Figs. 3c-d (∼ 20 mm deep in phantom). The two hypoechoic targets are not observable in the PA image due to no significant light absorption from the transparent agar-only medium, as expected.

**Figure 3:**
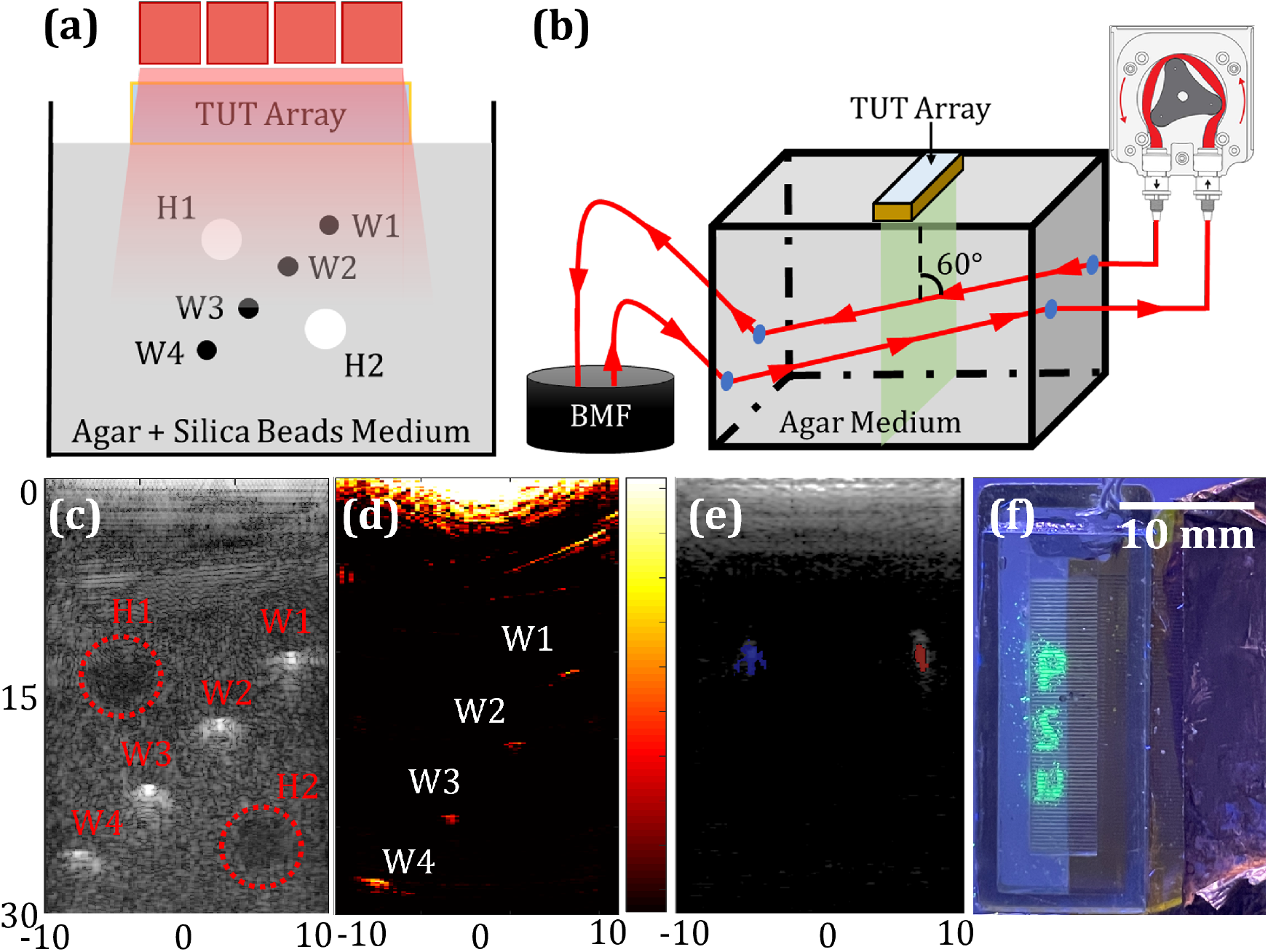
The TUT array based multimodal imaging validation. Phantom schematics for **(a)** ultrasound (US) and photoacoustic (PA) imaging and **(b)** Doppler ultrasound imaging. BMF: blood mimicking fluid. Imaging results showing **(c)** US image shows acoustic impedance contrast from the four micro metal wires (W1 -W4) and two hypoechoic targets (H1 and H2 in red dotted circles); **(d)** Photoacoustic image shows optical absorption contrast from the same four micro metal wires (W1-W4). Color bar on the right side shows the normalized PA amplitude in the range of 0 to 1; **(e)** The co-registered US and color Doppler US image of the phantom in (b) shows the ultrasound speckle contrast in gray scale and the Color Doppler US image in color scale. While the BMF flowing tubes showed the same US contrast, the opposite flow directions of blood mimicking fluid (BMF) in the two tubes are color coded in blue and red. Subfigures (c), (d), and (e) have axes units in mm. **(f)** The fluorescence image of the ultra-violet excitable beads patterned as letters ”PSU” imaged through the TUT array.

#### Doppler ultrasound

To demonstrate that the fabricated TUT array is sensitive for mapping the particle motion induced ultrasound frequency changes, Doppler ultrasound imaging was performed using a phantom consisted of a polyethylene tube running circulated blood mimicking fluid (BMF). The schematic of this phantom is shown in Fig. 3b and details are provided in Methods: Multimodal imaging phantom preparation. A peristaltic pump was used to circulate the BMF in a loop through the tube (hence two parallel tubes in the field of view), while the TUT array directly coupled to the phantom at an angle of 60°to the two tubes. The coregistered US and color Doppler image acquired and processed by Vantage 256 (see Methods: Imaging system and data acquisition sequence) in Fig. 3e, showed the same US speckle contrast from the two tubes in grayscale and an overlaid color Doppler image showed the opposite flow directions of the BMF in the two tubes in blue and red color scales. The measured size of these colored regions agreed well with the tube diameter of ∼2 mm.

#### Fluorescence imaging

In the final step, we validated the feasibility of fluorescence imaging through the TUT array. For this purpose, ultra-violet (UV) reactive fluorescence beads with 50 *µ*m beads diameter (Ultraglow, Techno Glow, Ennis, Texas, USA) were shaped to form ”PSU” pattern within a rectangular area of 12 mm × 3 mm. The The TUT-array was then placed on top of the pattern. Under the 120 Watts UV light excitation, the captured fluorescence emission signal from the ”PSU” could be easily distinguished from the background in Fig. 3f. The discontinuities in the fluorescence image is due to the translucent kerfs in the TUT array.

## 3 Discussions

Real-time multimodal imaging of optical and ultrasound contrasts of deep tissue using a single hand-held imaging device is ideally suited for several biomedical applications, such as endoscopy image guided biopsy or surgery. Conventional ultrasound imaging devices provide B-mode US based deep tissue structural information and Doppler US based functional blood flow information. However, they lack necessary molecular sensitivity for imaging the diseased tissue. On the other hand, optical contrast from both endogenous (melanin, hemoglobin, NADH) and FDA approved exogenous (e.g., indocyanine green) contrast agents have the capability to image the diseased tissue with high molecular sensitivity. While fluorescence imaging provides excellent molecular sensitivity, the PA imaging maps optical absorption contrast from deep tissue with high ultrasonic spatial resolution. US, fluorescence, and PA imaging technologies have been independently realized in many forms for different clinical applications, however, concurrent multimodal imaging requires effective integration of these imaging modalities into a single device. Because conventional US imaging devices are not transparent to light, integration of fluorescence and PA imaging to conventional US imaging devices is complicated and leads to shadow illumination problems.

To overcome the above problems, in this paper, a transparent lithium niobate based TUT-array was fabricated and validated for multimodal optical and ultrasound imaging applications. It is, to the best of our knowledge, the first transparent ultrasound linear array using bulk piezoelectric material. We successfully demonstrated the feasibility of the TUT array for a quad-mode US, PA, Doppler US, and fluorescence imaging in real-time. Dual-modality USPA imaging results using the TUT array demonstrated the potential of the TUT array to acquire both US (30 Hz frame rate) and PA (10 Hz frame rate, limited by laser firing rate) images with a bare minimum acoustic coupling between the array and the imaging subject. The ability to illuminate light through the TUT array and into the imaging object without any additional optical components not only reduced the complexity for building a multimodal ultrasound and optical imaging platform, but also helped eliminate the shadow illumination problems commonly observed in the conventional B-mode dual-modality USPA imaging systems. Further, the experiments on blood flow mimicking phantoms demonstrated that the TUT array is also capable of mapping the direction of blood flow using color Doppler ultrasound. In addition, the high optical transparency of the TUT arrays was exploited for imaging fluorescence objects. Together these experiments demonstrated the feasibility of developing multimodality optical and ultrasound imaging platform based on the TUT array technology, in particular the space constrained miniaturized multimodal endoscopy devices. For example, optical and ultrasound endoscopy technologies are the current standard for image-guided biopsy for cancer early detection of several internal organs [48–50], but patients suffer from multiple insertions due to current inability of combining optical and ultrasound biopsy technologies. Using the proposed multimodal TUT-array technology, the patient would only undergo one endoscopy procedure, which provides US based structural information, Doppler US based functional blood flow, PA based blood oxygenation information, and fluorescence enhanced tumor metabolism (e.g., NADH). Such a comprehensive information is needed to assess the tissue function and pre- and post-treatment efficiency [51].

However, the current TUT-array needs further optimization. The current TUT-array is suffering from limited bandwidths and sensitivity, which were mainly due to suboptimal transparent acoustic stacking materials: The glass bonded to the back side of the linear array induced the mass-loading effect and this resulted in a shift in the center frequency. Additionally, conductive glass slide is not an ideal backing material to the ultrasound transducer because its acoustic impedance (∼ 11 MRayls) is not matched to that of LN piezoelectric material (∼ 34 MRayls). In addition, glass slide is not a good acoustic absorbing or a damping material. These factors contributed to the broadening of the detected ultrasound spatial pulse length, and therefore reducing the bandwidth and deteriorating the axial resolution. There is a huge scope to further improve the TUT array bandwidth and sensitivity through various ways. The bandwidth can be primarily improved with better acoustic stacking. For example, the above mentioned glass as the first backing may be replaced by a transparent epoxy block coated with low temperature ITO deposition as a conductive connection for the ground to reduce the significant mass-loading effect from the glass slide. Further a two matching layer approach [52] or novel matching layers that use translucent particles (glass beads) also have the potential to help improve the transmit and receive bandwidths of the TUT arrays. For reducing the grating lobes, a geometry of fully diced through piezoelectric material would help reduce the cross-talk, which will likely reduce the grating lobes presented. The piezoelectric material may also be replaced with novel transparent piezoelectric materials, such as an alternative current (AC) poled PMN-PT [52, 53], which demonstrated higher piezoelectric coefficient than transparent LN and will therefore likely help enhance the dual-modality US and PA imaging sensitivity, imaging depth and spatial resolutions.

Despite these limitations, the new TUT array fabricated using transparent LN demonstrated potential advantages in realizing an integrated multimodal optical, US and PA imaging device for providing complementary structural, functional and molecular contrasts and spatial resolutions. The TUT platform can be scaled to develop multimodal devices of different length scales such as miniaturized endoscopy or wearable devices and therefore may open new avenues for combined US and optical imaging in pre-clinical and clinical studies.

## 4 Materials and Methods

### 4.1 TUT-Array fabrication and packaging

The TUT-array was fabricated by dicing a rectangular double-side polished transparent lithium niobate (LN) piece. The step-by-step fabrication process is illustrated in Fig. 4a. Step (1): a 0.5 mm thick 36°Y-cut LN wafer (Precision Micro Optics, Burlington, MA, USA) was used for the designed center frequency of 6.5 MHz. Step (2): 200 nm indium tin oxide (ITO) was deposited on both sides of the LN as a transparent and conductive electrode. Step (3): to form a common ground electrode, a one side ITO-coated glass slide was hard pressed to the ITO-coated LN using a small drop of transparent epoxy (EPO-TEK 301, Epoxy Technologies Inc., Billerica, MA, USA) as the bonding agency. Step (4): A high-precision dicing machine (K&S 982-6, Giorgio Technology sales/service, Mesa, AZ, USA) with a 70 *µ*m thick blade was used to dice out 64 elements on the LN substrate with 0.3 mm pitch. The dicing depth was kept to be ∼80% of the depth (400 *µ*m), so the ground electrode was intact. Step (5): A custom designed and fabricated polyimide-based flexible cable (70-pin with 0.3 mm pitch at the array-end and 0.5 mm pitch at the interface-end) was bonded to the transparent array by anisotropic conductive film (ACF) bonding. To maximize the transparent aperture and minimizes the acoustic mismatching effect on array elements, the polyimide flex cable was bonded to the elements with minimal overlap (∼2 mm). Step (6) Ground wires were connected to the ITO-coated glass plate electrode with help of a small blob of silver epoxy (E-solder 3022, Von Roll Isola Inc., New Haven, CT, USA). Step (7): for protection and electromagnetic shielding purpose, a rectangular brass housing (size 12.7 mm × 38.1 mm × 10 mm) was used to surround the device. A transparent epoxy (EPO-TEK 301, Epoxy Technologies Inc., Billerica, MA, USA) was then poured inside the brass housing to fill the kerfs and serve as the backing layer. A glass slide was placed on top of the brass tube to form a leveled epoxy layer. Step (8): Lastly, an 80 *µ*m thick Parylene C film was deposited on the full device to serve as the matching layer and waterproof layer. Matching layer can improve the acoustic wave transmission efficiency, and moreover, waterproof layer is critical to this device as ACF bonding tape is susceptible to humidity and can easily detach from the bonding. The fabricated linear array was then connected to a 70-pin 0.5 mm pitch commercial interface board (FPC050P070, Chip Quik Inc., Ancaster, ON, USA) by ACF bonding the interface end of the custom designed flexible circuit. Copper tape wrapped 42 AWG coaxial cables (2420/42 WH-100, Daburn Electronics & Cable, Dover, NJ, USA) were drawn from the interface board to Vantage 256 (Vantage 256, Verasonics Inc., Kirkland, WA, USA), a MATLAB based ultrasound data acquisition and image reconstruction platform, to perform the array characterization and US, PA and Doppler US imaging validations.

**Figure 4:**
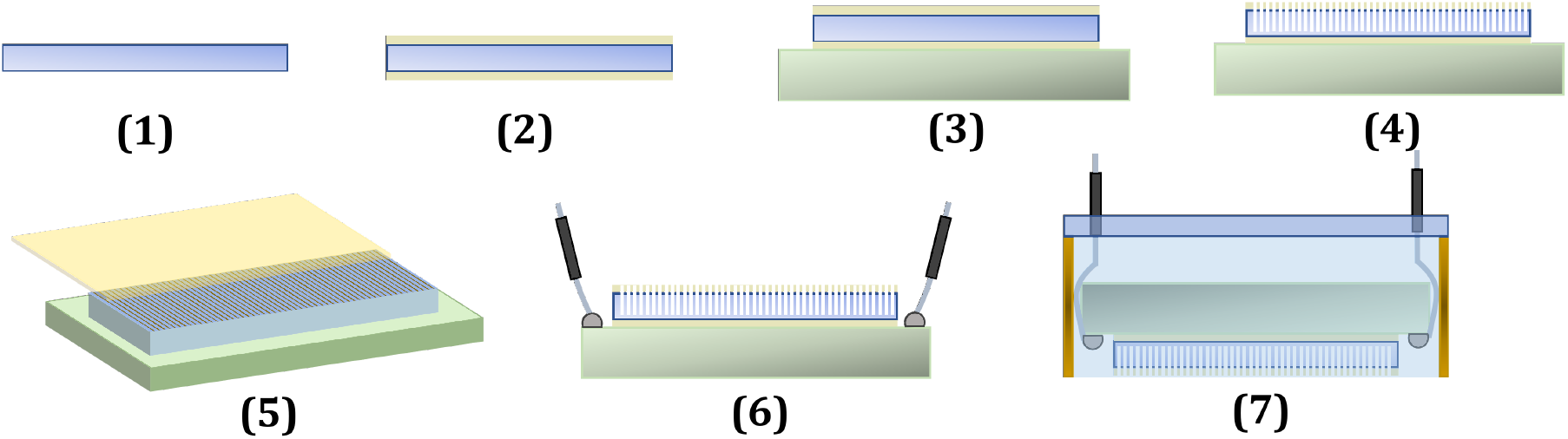
Step wise fabrication process of the TUT-array. **Step 1**: A double-side polished lithium niobate. **Step 2**: Indium tin oxide was deposited on two sides of the lithium niobate as conductive electrode. **Step 3**: The coated lithium niobate was bonded to a conductive glass slide. **Step 4**: the elements were created by dicing the lithium niobate to 80% depth. **Step 5**: the custom-made flexible cable was bonded to the array by anisotropic conductive film bonding. **Step 6**: The wires were connected to the conductive glass slide as ground. **Step 7**: the array was then placed inside a brass housing and filled with transparent epoxy

### 4.2 Imaging system and data acquisition sequence

#### USPA imaging

a portable optical parametric oscillator (OPO) laser source (Phocus Mobile, Opotek Inc., Carlsbad, CA, USA) was integrated with the Vantage system to provide 10 Hz, pulsed (5-7 ns pulses with peak output pulse energy of 120 mJ at 730 nm) and tunable (690-970 nm) optical illumination needed for multispectral PA imaging. A custom designed optical fiber bundle (FiberOptics Inc., CA, USA) that has a similar aperture area as the TUT-array aperture was employed to couple the light from the laser source to the array. Each B-mode US frame involved 64 focused transmit and receive acquisitions with only 8 transmitters active (where possible) on each side of the transmitting element and all 64 elements active for the US acquisition. Overall US frame rate achieved was 30 Hz. One laser pulse is fired each 100 ms and a corresponding PA image is generated with all 64 elements active in the receive mode. Thus, the PA imaging frame rate is 10 Hz, limited by the laser pulse repetition frequency. A function generator was used in master mode to synchronize both Vantage and the laser system, by setting the required time delays and thus allowing a proper interleaved, coregistered US + PA image formation.

#### Doppler ultrasound imaging

For acquiring the Doppler ensemble, 14 plane wave pulses were transmitted at an steering angle of 12° and at a pulse repetition rate of 3 kHz. The transmit pulses used for Doppler ultrasound acquisition consisted of three complete cycles in contrast to one cycle used for B-mode acquisition to get higher Doppler sensitivity. All 64 transmit and receive channels were active for each plane wave acquisition of the Doppler ensemble. The velocity and power Doppler processing was asynchronous with respect to the Doppler ensemble acquisition and was performed using the Doppler processing routines provided by the Verasonics platform.

### 4.3 Multi-modal imaging phantom preparation

#### USPA imaging phantom

four 50 µm diameter micro metal wires (W1-W4) were dyed using India ink to generate both ultrasound and photoacoustic contrasts. Micro metal wires were placed 5 mm apart from each other on the axial plane and 3 mm apart from each other on the lateral plane in an acrylic tank filled with a solution mixture of agar and silica beads. Silica beads and agar are mixed with water at 1% and 1.5% weight ratio respectively. Here the silica beads were used to mimic the background ultrasound speckle contrast. Then 1% agar solution was filled inside the 4.75 mm diameter cylindrical columns, next to the pencil leads, to serve as hypoechoic targets inside the tissue phantom for US imaging validation.

#### Blood flow Doppler phantom

A blood mimicking fluid (BMF-US, Shelley Automation, North York, Ontario, Canada) is circulated inside a polyethylene tube (outer diameter: 2.08 mm, inner diameter: 1.57 mm) using a peristaltic pump (model 3386, Cole-Parmer, Vernon Hills, IL, USA). The tube was partial submerged inside a tank filled with 1.5% agar solution. The tube was placed at 60 degrees to the imaging plane of the TUT-array as shown in Fig. 3b.

## Acknowledgments

We acknowledge Eugene Gerber for his kind support in the machining of parts and help with transducer design. We also thank Christopher Cheng with his help in flexible cable fabrication and Parylene coating.

## Author Contributions

Conceptualization, H.C., A.D., M.O., and S.-R.K.; software, H.C., S.M., and S.A.; investigation, T.J. and S.-R.K.; data curation, H.C., M.O., and S.A.; writing—original-draft preparation, H.C.; experiment design, H.C., and S.-R.K.; hardware, H.C., A.D., J.M., J.L., Q.T., and M.O.; supervision, S.-R.K.; project administration, S.-R.K.; funding acquisition, S.-R.K

## Data Availability

Data underlying the results presented in this paper are not publicly available at this time but may be obtained from the authors upon reasonable request.

